# VIT1-dependent Fe distribution in seeds is conserved in dicots

**DOI:** 10.1101/503342

**Authors:** Seckin Eroglu, Nur Karaca, Katarina Vogel-Mikus, Anja Kavčič, Ertugrul Filiz, Bahattin Tanyolac

**Affiliations:** Department of Genetics and Bioengineering, Izmir University of Economics, Izmir, Turkey; Department of Bioengineering, Ege University, Izmir, Turkey; Department of Biology, University of Ljubljana, Ljubljana, Slovenia; Jozef Stefan Institute, Jamova 39, 1000 Ljubljana, Slovenia; Cilimli Vocational School, Department of Crop and Animal Production, Duzce University, Duzce, Turkey

**Keywords:** Biofortification, seed, iron, metal, vit1, plastid, synchrotron, homeostasis

## Abstract

One third of the people suffer from iron (Fe) Fe deficiency. An underlying factor for this malnutrition is insufficient Fe intake from the diet. A major part of the human diet includes seeds of staple crops, which contain Fe that is poorly bioavailable. One reason for the low bioavailability is these seeds store Fe in cellular compartments that also contain antinutrients, such as phytate. Thus, several studies focused on decreasing phytate concentrations. As an alternative approach to increase bioavailable Fe, Fe reserves might be directed to cellular compartments such as plastids that are free of phytate. Previous studies indicated that Fe reserves can be relocalized inside the seed to the desired compartment by genetic modification, provided that a suitable iron transporter protein is used. However, to the best of our knowledge, a Fe transporter localizing to plastids have not been identified in seeds to date. To discover novel Fe transporters, we screened Fe patterns in seeds of distinct plant lineages, hypothesizing Fe hyperaccumulating sites would indicate Fe transporter presence. To this end, metal localizations in seeds of more than twenty species were investigated using histochemical or X-ray based techniques. Results showed that in Rosids, the largest clade of eudicots, Fe reserves were primarily confined in the embryo part of the seeds. Furthermore, inside the embryos, Fe was enriched in the endodermal cell layer, a well-known feature that is mediated by vacuolar Fe transporter, VIT1 in model plant *Arabidopsis thaliana*. This enrichment was well conserved in and beyond Rosid species. Finally, a few seeds showed novel Fe patterns, including *Carica papaya* which concentrated large Fe reserves exclusively in plastids called amyloplasts. Generally, Fe stored in amyloplast is considered bioavailable. Taken together, this study suggests dicot seeds store Fe mainly in the embryo, with a VIT1-dependent enrichment in its endodermal cell layer and indicate *Carica papaya* possess a strong Fe transporter at the plastid membrane. Once it is identified that might be useful in biofortification, as a novel tool to shift Fe to compartments where it is more bioavailable.

## Introduction

One-third of the people suffer from Fe deficiency (WHO, 2014). One of the main reasons for this malnutrition is insufficient Fe intake from the diet. Human diets are largely based on plants which might be a poor source of dietary Fe (Gibson et al., 2010). This is true especially when edible parts of the major staple crops, seeds are considered. Corn, wheat, rice kernels not only contain low concentrations of total iron but also, large part of it cannot be absorbed by the human digestive system (Borg et al., 2009). To combat Fe deficiency, modifying plants to fortify seeds with more Fe, also called biofortification of seeds with Fe, is proposed to be the most sustainable approach (Shahzad et al., 2014).

Genetic engineering, classical breeding and other methods have been applied to increase Fe concentration in the seeds (Murgia et al., 2012). The classical breeding has resulted in limited success especially when natural cultivars show low variability in seed Fe levels. Genetic engineering methods successfully increased Fe levels in major crops such as rice and wheat (Connorton et al., 2017; Lee et al., 2009). This increase has been mainly achieved by expressing proteins that boosts Fe storage capacity of the cells, eventually forcing mother plant to divert more Fe into the seeds. Most of the Fe biofortification approach aimed at increasing Fe concentration; however, not only the concentration of Fe but also its bioavailability is a major concern.

Bioavailability determines the portion of the Fe that can actually be absorbed by the human digestive system. This portion can be less than ten percent of the total iron, but it is strictly dependent on the form of Fe (Hurrell and Egli, 2010). In staples Fe is accumulated in cells that contain metal binding molecules, phytate being the most important. Fe is released from phytate by the plant itself during germination without a hassle due to the presence of phytases (Hegeman and Grabau, 2001). However, humans do not possess phytase, thereby phytates act as antinutrients in humans. To increase Fe bioavailability, several studies aimed at decreasing phytate concentrations (Reddy et al., 2017; Shi et al., 2007). Such a decrease has been achieved, but many times at the expense of severe growth problems in the plant (Warkentin et al., 2012). An alternative approach could be to direct Fe from phytate containing compartments to phytate-free compartments since antinutrients are unevenly distributed in cells. In nature, in subcellular level, Fe can be stored in vacuoles, plastids, cytosol or elsewhere (Cvitanich et al., 2010; Otegui et al., 2002). In vacuoles, Fe can be found in phytate-rich compartments called globules (Davila-Hicks et al., 2004; Lanquar et al., 2005). In plastids, Fe is associated with ferritin protein (Briat et al., 2010; Waldo et al., 1995). In contrast to antinutrient bound Fe, ferritin bound Fe is bioavailable (Davila-Hicks et al., 2004). Thus, optimizing seed Fe for human nutrition requires not only increasing Fe concentration but also its bioavailability, which is strictly dependent on its subcellular level distribution.

The advances in Fe imaging techniques allowed detailed investigation of Fe localization in small seeds of *Arabidopsis thaliana*, which eventually led to discovery of Fe storage proteins. X-ray synchrotron fluorescence imaging technique has been used for the first time in plant science to identify that Fe is not homogeneously distributed throughout the embryo but rather accumulated around the provascular strands (Kim et al., 2006). However, the low resolution of the technique at that time did not allow to pinpoint the specific cells that are overaccumulating Fe. Such a resolution has been achieved with the use of an enhanced version of classical Perls staining; Perls/DAB, namely (Roschzttardtz et al., 2009). Perls/DAB revealed that the Fe enrichment around provascular strands is confined to vacuoles of endodermal cells (Ramos et al., 2013; Roschzttardtz et al., 2009).

By comparison of Fe localizations in embryos of T-DNA insertion mutants of *Arabidopsis thaliana*, the genetic basis of Fe transport into endodermis has been tackled. Two independent lines that have mutations in V*ACUOLAR IRON TRANSPORTER1 (VIT1)* gene, completely lost the Fe enrichment in their endodermis (Eroglu et al., 2017; Kim et al., 2006). *VIT1* localized to the tonoplast, expressed in the stele, and complemented Fe hypersensitivity of yeast when expressed heterologously; thereby proposed to be a vacuolar Fe importer in endodermis (Kim et al., 2006). Disruption of VIT1 results in relocalization of Fe reserves in a distinct Fe hotspot which is mediated by METAL TOLERANCE PROTEIN8 (MTP8) (Eroglu et al., 2017). Finally, in *vit1 mtp8* double knock out, Fe relocalizes homogeneously. In conclusion, analysis of loss-of-function mutants of *Arabidopsis thaliana* shows that metal transporters determine where metals will be stored in the mature seed. This indicates that if a suitable Fe transporter protein is used, directing Fe to certain organelles that are free of antinutrients might be possible.

Our research aim was to identify a seed Fe transporter that can be useful to direct Fe to the plastids. *Arabidopsis thaliana*, accumulating Fe almost exclusively inside the vacuoles, was not suitable for such an approach. Whereas, Fe distribution in seeds of other plants is scarcely known. To find plastid targeting Fe transporters, we screened Fe distributions in seeds belonging to distinct plant lineages. Based on what is known from the model plant, we hypothesized that Fe concentrated regions will indicate the presence of a transporter. To this end, seeds of more than 20 different species belonging to several different plant orders were examined by visualizing their Fe patterns by using Perls/DAB and synchrotron X-ray fluorescence spectroscopy. To our surprise, results show that the most common Fe enriched region corresponded to endodermis, a well-known Fe enriched region mediated by VIT1 gene in *Arabidopsis thaliana*. Since VIT1 was discovered, it has attracted a wide attention from seed Fe research; nevertheless, whether it is conserved in other plant species or not, remained elusive. Therefore, we also investigated how widespread the Fe enrichment in the endodermis, in addition to looking for new Fe hotspots.

## Material and methods

### Collection of seeds

Seeds were obtained from gene banks, local suppliers or personally collected. These are listed in Table 1.

**Table 1:** Seeds used in the study. Species

| Species | Source |
| --- | --- |
| *Linum usitatiss* | National Genebank, Menemen, Turkey |
| *Medicago truncatula* | National Genebank, Menemen, Turkey |
| *Cucumis sativus* | National Genebank, Menemen, Turkey |
| *Citrullus lanatus* | National Genebank, Menemen, Turkey |
| *Cannabis sativa* | National Genebank, Menemen, Turkey |
| *Euonymus europaeus* | National Genebank, Menemen, Turkey |
| *Alyssum sibiricum* | National Genebank, Menemen, Turkey |
| *Brassica oleracea* | National Genebank, Menemen, Turkey |
| *Gossypium arboreum* | National Genebank, Menemen, Turkey |
| *Batis Maritima* | Kew Gardens, London, England |
| *Moringa peregrina* | Kew Gardens, London, England |
| *Eucalyptus elata* | Kew Gardens, London, England |
| *Geranium carolinianum* | Kew Gardens, London, England |
| *Brassica napus* | Genebank, Gatersleben, Germany |
| *Limnanthes douglasii* | Botanischer Garten, Marburg, Germany |
| *Arabidopsis thaliana* | Arabidopsis Biological Resource Center,<br>Ohio, USA |
| *Reseda lutea* | Personally collected* |
| *Noccaea praecox* | Personally collected** |
| *Carica papaya* | Commercial |
| *Arachis hypogaea* | Commercial |
| *Solanum lycopersicum* | Commercial |
| *Capparis spinosa* | Commercial |

\*(Çiğdem et al., 2018). \*\*collected from Pb, Zn polluted site, Mežica, Slovenia (Vogel- Mikus et al., 2005)

### Perls staining and DAB/H_2_O_2_ intensification (Perls/DAB)

Wherever possible (embryos were large, could easily be isolated, seed coat was sufficiently soft to be cut, and high magnification was not needed) samples were directly (i.,e., without fixation) stained by Perls (Fig. 4 A, B) or Perls/DAB (Fig. 2C, 3D) (Roschzttardtz et al., 2009). In other cases, *in situ* protocol was used. For direct Perls staining, samples were incubated in 4% (v/v) HCl and 4% (w/v) K-ferrocyanide (Perls stain solution) for 15 min and incubated for 30 min at room temperature (Stacey et al., 2008). If the Fe signal was weak, it was intensified using DAB (Roschzttardtz et al., 2009). Perls stained samples were incubated in a methanol solution containing 0.01 M NaN_3_ and 0.3% (v/v) H_2_O_2_ for 1 h, and then washed with 0.1 M phosphate buffer (pH 7.4). For the intensification reaction samples were incubated between 10 to 30 min in a 0.1M phosphate buffer (pH 7.4) solution containing 0.025% (w/v) DAB, 0.005% (v/v) H_2_O_2_, and 0.005% (w/v) CoCl_2_. The reaction was stopped by rinsing with distilled water.

**Figure 3:**
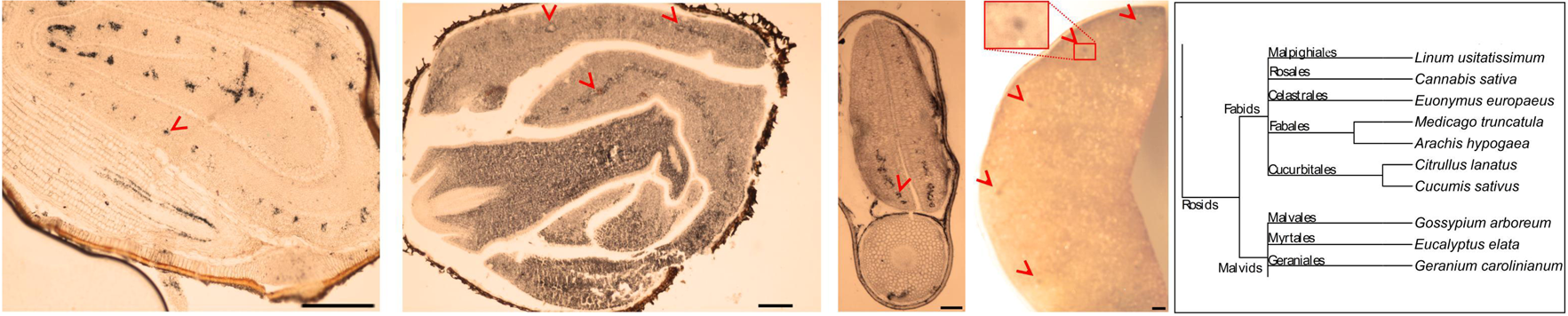
Specific Fe enrichment in endodermis is even conserved in distinct plant orders. Seeds belonging to various orders were cut and thin cross sections were obtained. Cross sections were stained with Perls/DAB and observed under light microscope. From **(A-C)**, *Gossypium arboreum*. *Eucalyptus elata* and *Medicago truncatula*, respectively. **(D)** single cotyledon of *Arachis hypogaea*. Fe staining appeared black in **(A-C)** and brown in **(D)**. Outer brown-black cover in **(A-C)** are seed coats. These were already brown before the staining, thus the color do not reflect the stained Fe. In **(A)**, longitudinal section of hypocotyl and cross section of folded cotyledons can be differentiated. Red arrow heads show Fe enrichment surrounding the provascular strands of cotyledons. **(E)** a branch of the taxonomic tree, refer to Fig. 8 for the whole tree. This branch shows species that were examined for Fe reserves in distinct orders and used as a visual aid. Note that only selected examples from **(E)** is shown through **(A-D)**. Bar is 1 mm for **(A)** and 0.1 mm for **(B-D)**. Red arrow heads point to examples of specific Fe accumulation pattern, closed rings around provasculature of cotyledons.

**Figure 4:**
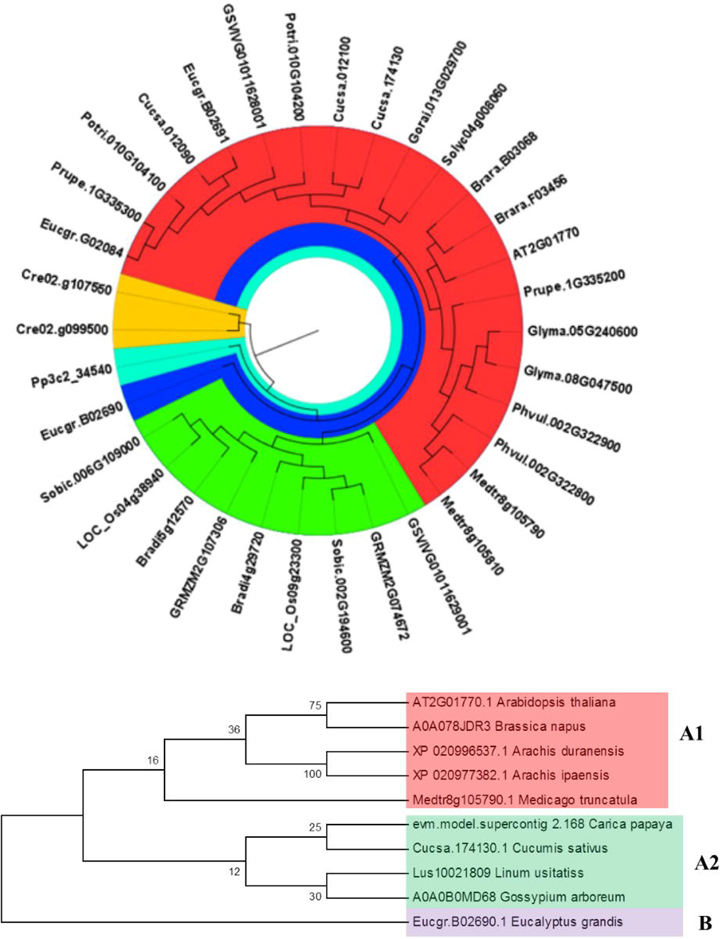
Phylogenetic distribution of VIT1 orthologs in 18 different plant species from monocots, dicots and lower plants. **(A)** phylogenetic distribution of VIT1 orthologs in 18 different plant species from monocots, dicots and lower plants. Phylogeny was constructed by MEGA 7 with ML method for 1000 bootstraps using putative 34 VIT1 protein sequences. The red segment includes only dicots. Green segment includes only monocots except for one member of *Vitis vinifera*. The blue segment shows a diverged dicot member of *Eucalyptus grandis*. Orange and cyan colors show algal and moss species, respectively. For details on analyzed sequences, refer to materials and method section. **(B)** a phylogeny was constructed by MEGA 7 with ML method for 1000 bootstraps.

For *in situ* Perls staining, seeds were first fixed in 10% formalin and then dehydrated in an ethanol series (30%, 40%, …, 100%). Seeds were cleared with xylene. Seeds that were still too hard to cut were further incubated in 20% formic acid for 45 minutes. Seeds were then embedded in wax and 10-30 µm sections were cut and stained (Roschzttardtz et al., 2009). Sections were treated with 4% (v/v) HCl and 4% (w/v) K-ferrocyanide (Perls stain solution) for 15 min and incubated for 30 min at room temperature (Stacey et al., 2008). DAB intensification was applied as described in Meguro et al. (Meguro et al., 2007). For the intensification reaction the sections were incubated between 10 and 30 min in a 0.1 M phosphate buffer (pH 7.4) solution containing 0.025% (w/v) DAB (Sigma), 0.005% (v/v) H_2_O_2_, and 0.005% (w/v) CoCl_2_ (intensification solution). The reaction was stopped by rinsing samples with distilled water. Samples were imaged with a light microscope (Axioskop; Carl Zeiss, Jena, Germany).

### X-ray fluorescence

X-ray fluorescence imaging of *Brassica napus, Moringa peregrine* and *Euonymus europaeus* was performed at XRF beamline of Synchrotron Elettra (Karydas et al., 2018) on 60 µm seed cross-sections. The seeds were imbibed overnight at 4°C, flush frozen in propane cooled with liquid nitrogen, embedded in tissue freezing medium (Leica, Germany) and cut using CM3060 Leica Cryostat(Vogel-Mikuš et al., 2014). The sections were sandwiched between two 2.5 µm Mylar foil and scanned by 75 × 75 µm beam at an excitation energy of 10 KeV. Obtained XRF spectra were fitted by PyMCA software (Solé et al., 2007) and quantified (Kump and Vogel-Mikuš, 2018). Imaging of element distribution in *Noccaea praecox* was performed by micro-PIXE as described (Vogel-Mikus et al., 2007; Vogel-Mikuš et al., 2008, 2014).

### Bioinformatics Analyses

VIT1 protein sequences were derived from the Phytozome database 12.1.6 version (https://phytozome.jgi.doe.gov/pz/portal.html#; (Goodstein et al., 2011)) for bioinformatics analyses. Following plants were included: *Arabidopsis thaliana* (AT2G01770), *Carica papaya* (evm.model.supercontig_2.168), *Cucumis sativus* (Cucsa.174130.1, Cucsa.012090, and Cucsa.012100), *Linum usitatiss* (Lus10021809), *Medicago truncatula* (Medtr8g105790 and Medtr8g105810), *Eucalyptus grandis* (Eucgr.B02690, Eucgr.B02691, and Eucgr.G02084), *Brassica napus* (A0A078JDR3), *Gossypium arboreum* (A0A0B0MD68), *Arachis duranensis* (XP_020996537.1), *Arachis ipaensis* (XP_020977382.1), *Brachypodium distachyon* (Bradi4g29720 and Bradi5g12570), *Brassica rapa* (Brara.B03068 and Brara.F03456), *Chlamydomonas reinhardtii* (Cre02.g099500 and Cre02.g107550), *Glycine max* (Glyma.05G24060 and Glyma.08G047500), *Gossypium raimondii* (Gorai.013G029700), *Zea mays* (GRMZM2G074672 and GRMZM2G107306), *Vitis vinifera* (GSVIVG01011628001 and GSVIVG01011629001), *Oryza sativa* (LOC_Os04g38940 and LOC_Os09g23300), *Phaseolus vulgaris* (Phvul.002G322800 and Phvul.002G322900), *Populus trichocarpa* (Potri.010G104100 and Potri.010G104200), *Physcomitrella patens* (Pp3c2_34540), *Prunus persica* (Prupe.1G335200 and Prupe.1G335300), *Sorghum bicolor* (Sobic.002G194600 and Sobic.006G109000), and *Solanum lycopersicum* (Solyc04g008060). Protein domains were detected by using Pfam 31.0 (https://pfam.xfam.org; (Finn et al., 2015)). Sequence length, molecular weight, and isoelectric point (*pI*) were determined by using ExPASy ProtParam tool (https://web.expasy.org/protparam/, (Gasteiger et al., 2005)). The transmembrane helices were predicted by using The HMMTOP transmembrane topology prediction server version 2.0 (http://www.enzim.hu/hmmtop/, (Tusnady and Simon, 2001)). Phylogenetic tree of VIT1s was constructed with MEGA 7.0.2 software (Kumar et al., 2016) using maximum likelihood (ML) method with 1000 bootstraps. The evolutionary distances were computed using the Poisson correction method (Zuckerkandl and Pauling, 1965). Identity values (%) of VIT1s were analyzed through NCBI blastp tool (https://blast.ncbi.nlm.nih.gov/Blast.cgi?PAGE=Proteins).

## Results

### Most of the seed Fe reside in endodermis in all tested members of Brassicaceae family

To investigate how Fe distribution differs between *Arabidopsis thaliana* and other members of Brassicaceae, by using Perls/DAB method and X-ray synchrotron analysis, Fe distribution patterns in several different Brassicaceae species were visualized. *Arabidopsis thaliana* showed closed circle shaped Fe enriched regions in cotyledons and hypocotyl (Figure 1A), confirming the previous reports (Roschzttardtz et al., 2009). This region corresponds to endodermis (Roschzttardtz et al., 2009). Similar to *Arabidopsis thaliana* (Fig.1A), all other members of the Brassicaceae showed closed circular-like Fe enriched regions, both in cotyledons and hypocotyl (Fig.1B, C, D, E). Seemingly, Fe rings overlap a region surrounding provascular bundles, most probably endodermis, based on extrapolation from *Arabidopsis thaliana*. *Brassica napus* and *Brassica oleracea* contained conduplicate cotyledons (I.e., inner and outer), both of which exhibited the same Fe distribution pattern (Fig.1B, D). Despite the similarities in these patterns, variations between species were also observed. In contrast to *Arabidopsis thaliana* (Fig.1A), Fe enriched region of *Brassica napus* was not confined to a single cell layer in the hypocotyl (Fig.1C). Furthermore, *Alyssum sibiricum* showed two adjacent Fe enriched circles instead of one in its hypocotyl (Fig.1D). Taken together, results showed that not only *A. thaliana*, but also other Brassicaceae species store main Fe reserves in endodermis.

**Figure 1:**
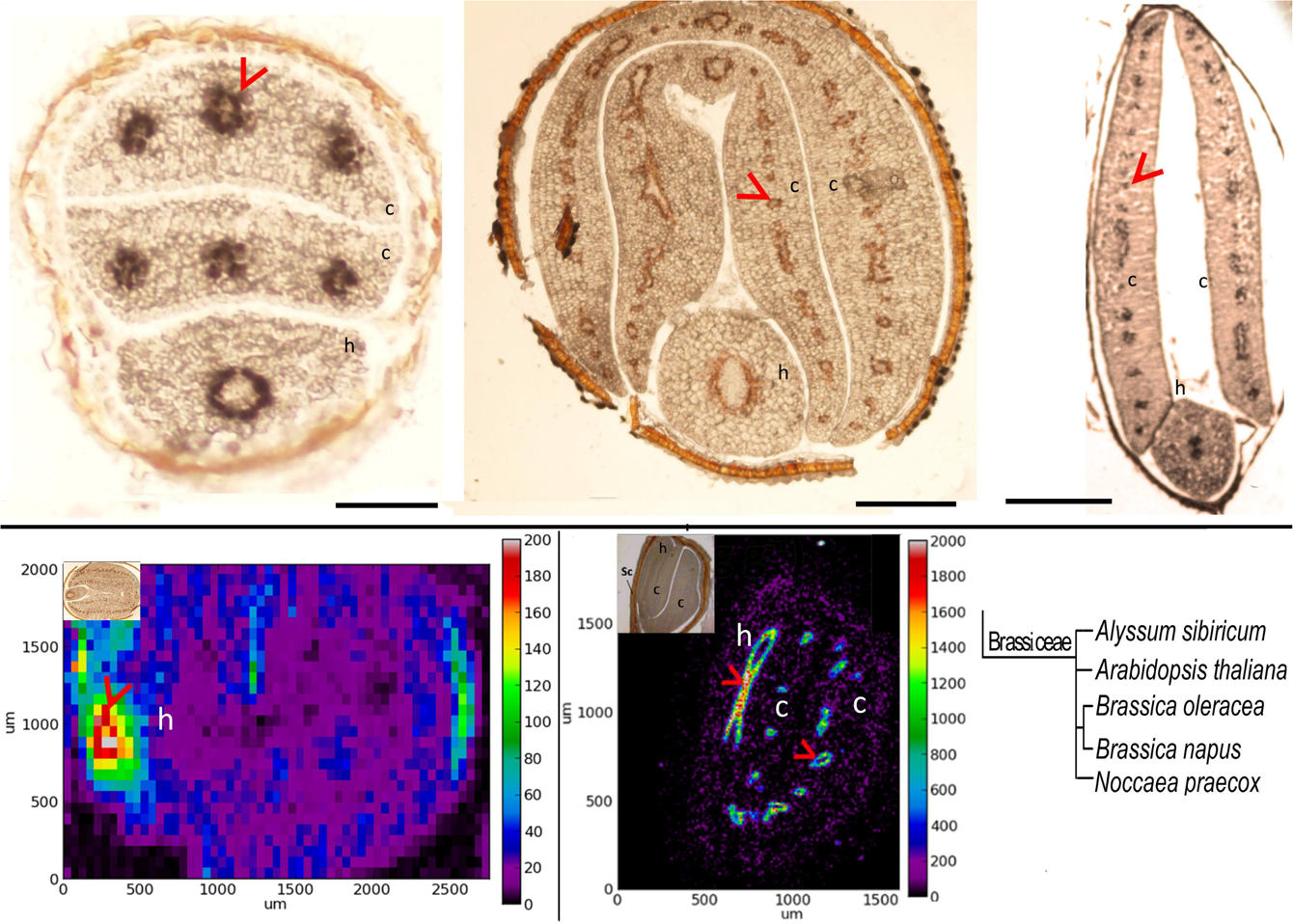
Fe is enriched in the endodermis in all examined species of Brassiceae family. **(A-C)** Perls/DAB stained seed cross sections. Seeds belonging to different members of Brassiceae family were cut and thin cross sections were obtained. Cross sections were stained with Perls/DAB and observed under light microscope. From left to right: *Arabidopsis thaliana*, *Brassica oleracea and Alyssum sibiricum*. Brown regions surrounding the embryos are seed coats. These were already brown before the staining; thus, the color does not reflect the stained Fe. Fe appeared as black stains on **(A)** and **(C)**, and brown in **(B)**. Circular-shaped staining in *Arabidopsis thaliana* corresponds to the endodermal cells surrounding the provascular strands. **(D, E)** Synchrotron X-Ray fluorescence images of relative Fe distribution in the seeds. **(D)** *Brassica napus* and **(E)** *Noccaea praecox.* **(F)** a branch of the taxonomic tree, refer to Fig. 8 for the whole tree. This branch shows species that were examined for Fe reserves in Brassiceae family and used as a visual aid. Note that, in contrast to *Arabidopsis thaliana*, *Alyssum sibiricum* and *Noccaea praecox*, which have a single pair of cotyledons, both *Brassica napus* and *Brassica oleracea* consist of a pair of inner and an outer cotyledon. Bar represents 0.1 mm in **(A)**, 0.5 mm in **(B-C)**. c: cotyledon, h: hypocotyl. Red arrow heads point to examples of specific Fe accumulation pattern, closed rings around provasculature of cotyledons **(A, B, C, E)** and hypocotyl **(D)**.

### Fe-enriched endodermis is conserved in the order Brassiccales

Next, whether conserved Fe pattern in *Arabidopsis thaliana* seed extends beyond the family level to the order level was assessed. Ring-like Fe distribution patterns around provasculature of either cotyledon or hypocotyl, which is the typical pattern that was seen in Brassicaceae were sought. Species of the several distinct families under the Brassicales were stained by either Perls alone (for Fe rich samples) or with DAB intensification (for those that produce low signal, i.e., low Fe concentration) to reach a balance in staining intensity. In *Limnanthes douglasii* embryos, at the first glance, not only endodermal cells but most of the others were stained (Figure 2A). However, closer examination around provascular bundles of cotyledons and comparison of these cells with nearby cells revealed that single cell layer around the strands (i.e., endodermis) were slightly enriched with Fe (Figure 2B). In *Capparis spinosa*, Perls staining without DAB amplification revealed that Fe was accumulated close to the central cylinder of the hypocotyl, most probably endodermis, based on the typical ring-like distribution of Fe in endodermis of other species (Fig. 2C). In *Batis Maritima*, Fe was stored in several cell layers surrounding the provasculature including endodermis (Fig. 2E). Therefore, Fe enriched endodermis that was initially observed in *A. thaliana*, is conserved in plants at least in order level.

**Figure 2:**
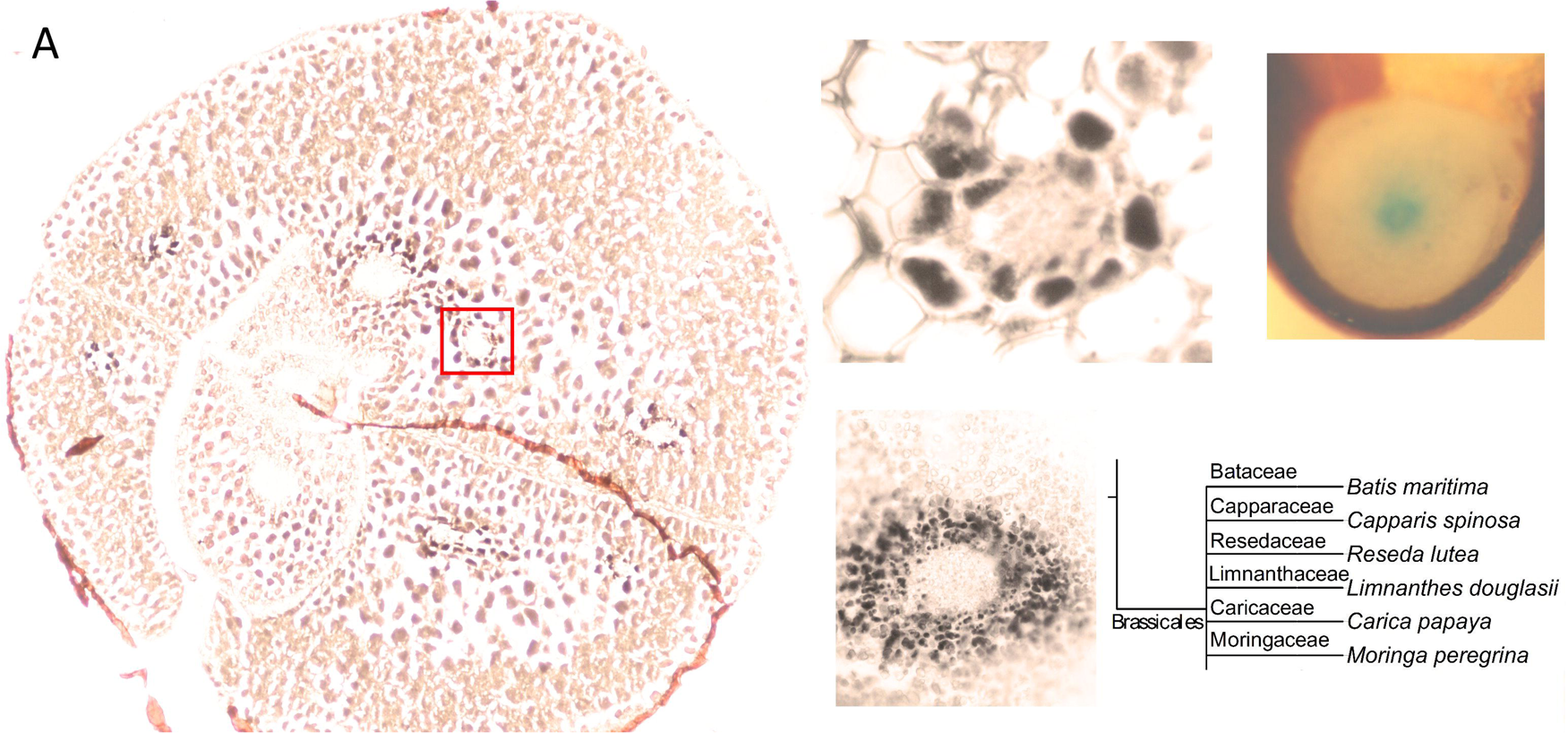
Examples of Fe enrichment in the endodermis is conserved in Brassicales order. Seeds belonging to different members of Brassiceae family were cut and thin cross sections were obtained. Cross sections were stained either with Perls or with Perls/DAB and observed under light microscope. **(A)** *Limnanthes douglasii*. **(B)** Higher magnification of the red square in **(A)**. **(C)** hypocotyl of *Capparis spinosa*. **(D)** hypocotyl of *Batis maritima*. **(B-D)** focusses on vascular tissues. Fe staining appeared black in **(A, B, D)** and blue in **(C)**. Brown colored structure partially surrounding the embryo in **(A)** is the seed coat. This was already brown before the staining; thus, the color does not reflect the stained Fe. This branch shows species that were examined for Fe reserves in Brassiceae family and used as a visual aid. Bar is 0.1 mm for **(A)**, **(C)**; 0.01 mm for **(B)** and **(D)**. **(E)**, branch of the taxonomic tree, refer to Fig. 8 for the whole tree. This branch shows species that were examined for Fe reserves in the same order but different families and used as a visual aid. Red arrow heads point to examples of specific Fe accumulation pattern, closed rings around provasculature of cotyledons **(A, B)** and hypocotyl **(C, D)**.

### Species other than the ones in Brassicales also overaccumulate Fe in the endodermis, although this often does not represent the largest Fe pool

Next step was to further pursue the Fe distribution pattern in orders distinct from Brassicales. Brassicales and sixteen other orders together constitute Rosids (Chase et al., 2016). Random species belonging to diverse orders under Rosids were collected (Table 1). *Gossypium arboreum* showed closed circular ring-like Fe distribution (Fig.3A), similar to typical Fe stained endodermal cells. Furthermore, endodermis represented the main Fe reserves in the seed. Likewise, *Eucalyptus elata* also showed Fe enrichment in the endodermis. However, this enrichment was evident in cotyledons but failed to be discerned in the hypocotyl, where cortical cells were already heavily stained (Fig.3B). Two species that belong to the Fagales order, *Medicago truncatula* and *Arachis hypogaea*, were examined. (Fig.3C, D) Fe staining in endodermis of *Medicago truncatula* was confined to the cotyledons and apparently represented the largest Fe pool in the embryo (Fig.3C). In contrast to *Medicago truncatula*, main Fe reserves in *Arachis hypogaea* were mostly homogeneously distributed. In a detailed examination, although endodermal cells of *Arachis hypogaea* did not represent the major Fe pool, they were still slightly enriched with Fe in comparison to cortical cells (Fig.3D).

### Conservation of VIT1 sequence in species that do not show an Fe enriched endodermis

In *Arabidopsis thaliana* seed, Fe enrichment in endodermis is strictly dependent on a functional VIT1 protein (Kim et al., 2006). We failed to observe Fe enriched endodermis in some species, indicating they may not have a functional VIT1. In order to test how well VIT1 sequence is conserved in plants, VIT1 sequences in distinct plants species were compared (Fig. 4). First, to get a general picture of how VIT1 is conserved in plants, 34 VIT1 sequences from 18 plant species were determined and compared (Fig.4A). In contrast to *Arabidopsis thaliana* that had only one VIT1, most plants showed two copies of it. Interestingly, in *Vitis vinifera*, one of the two copies of VIT1 was placed with monocot VITs in the phylogenetic tree. *Eucalyptus grandis* contained three homologs of VIT1, one of which was distinctly diverged. These data suggest that VIT1 is well conserved in distinct plant lineages (Supp. Fig. 1, 2).

Next, we assessed conservation of VIT1 protein sequence in species that were used in Fe staining. The whole genome sequence was available only for *Arabidopsis thaliana*, *Carica papaya*, *Cucumis sativus*, *Linum usitatiss*, *Medicago truncatula*, *Eucalyptus grandis*, *Brassica napus*, *Gossypium arboreum*, *Arachis sp.* (Fig. 4B). VIT1 sequences of all ten species formed three subgroups. The lengths of VIT1 sequences ranged from 245 to 265 amino acids with five transmembrane helices (Supp. Table 1). In addition, these contained VIT1 domain structure (PF01988) (Supp. Fig. 2) and indicated an acidic character. Protein BLAST analyses showed that identity values (%) *Arabidopsis* and other nine VIT1s ranged from 29% to 92% (Supp. Fig. 3). While the highest identity value was found as 92% between *Arabidopsis* and *Brassica napus*, followed by *Medicago truncatula* and *Linum usitasiss* as 83%, the lowest value was found as 29% between *Arabidopsis thaliana* and *Arachis* spp., indicating variations of *VIT1* genes in plants. All three groups possessed members that stored Fe around the provasculature. Therefore, although seeds of *Carica papaya* and *Linum usitatiss* did not exhibit a VIT1-dependent Fe enrichment in the endodermis (Fig. 5 and Supp. Fig. 4), their VIT1 sequences did not diverge remarkably from others.

**Figure 5:**
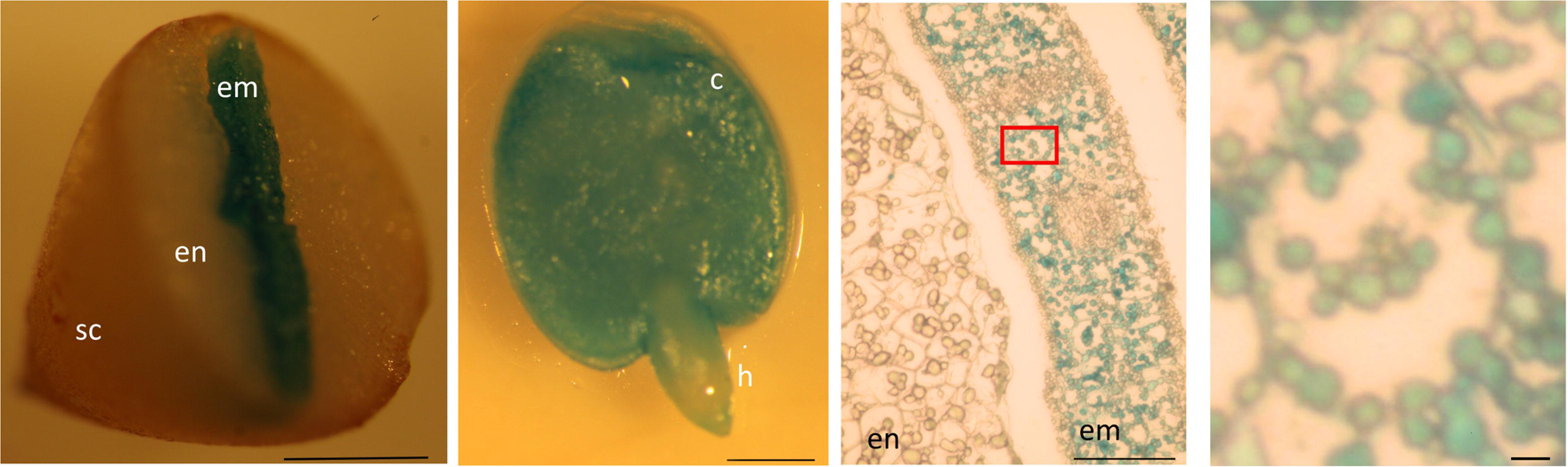
Supra- and sub-cellular level Fe storage in *Carica papaya* differs drastically from other stained seeds. Distribution of Fe in *Carica papaya* seeds. *Carica papaya* seeds were cleaned from the pericarp. **(A)** seeds were cut into half by hand and were stained by Perls. **(B)** whole embryo was isolated from the seed and was stained by perls. **(C)** the thin cross section of a whole seed, stained by Perls. **(D)** close-up to the red rectangle in **(C)**. Blue colour in all pictures indicate stained Fe. Blue small spheres resemble leucoplasts. sc: seed coat, en: endosperm, c: cotyledon, h: hypocotyl em: embryo. Bar is 1 mm for **(A, B)**; 0.1 mm for **(C)**; 0.01 mm for **(D)**.

### Seed Fe storage shows diversity in organ, tissue and subcellular level

Perls and Perls/DAB staining of seeds also revealed distinct ways of Fe accumulation. A hand-cut *Carica papaya* seed showed intense staining in the center, corresponded to the embryo (Fig. 5A). In contrast to many other Rosids, *Carica papaya* seeds contained a large endosperm tissue; however Fe reserves confined almost exclusively to the embryo (Fig. 5A, B). In order to further investigate Fe localization in tissue and subcellular level, *Carica papaya* seeds were fixed, cut and stained. They showed intense and clear-cut staining even the DAB intensification was skipped (Fig. 5C). Staining was confined to multiple tiny organelles similar to leucoplasts regarding the shape, size and number. Distribution of this organelle did not differ between endodermis and other cell layers. Therefore, although *Carica papaya* belongs to the same order with *Arabidopsis thaliana*, differs strikingly from it by storing Fe in homogeneously distributed leucoplasts throughout the embryo.

Nutrients are usually stored in embryos in Rosids. Besides embryo, endosperm accumulated Fe significantly in a few Rosid species. *Euonymus europaeus* contained a large endosperm which was stained by Perls/DAB heavily (Fig. 6A-B). In addition to the endosperm, we found examples where seed coat stored significant Fe. Usually, colourful seed coats prevent detection of Perls/DAB staining. However, *Moringa peregrina*, which contains two thick layers of seed coat, lacks a color in its inner seed coat (Muhl et al., 2016). Therefore, not only *Moringa peregrina’s* embryo but also its seed coat was stained by Perls/DAB (Fig. 6C-D). Interestingly, the intensity of staining was higher in its seed coat compared to that of the embryo (Fig. 6C). This indicated that a major Fe accumulation site in *Moringa peregrina* was found in the seed coat.

**Figure 6:**
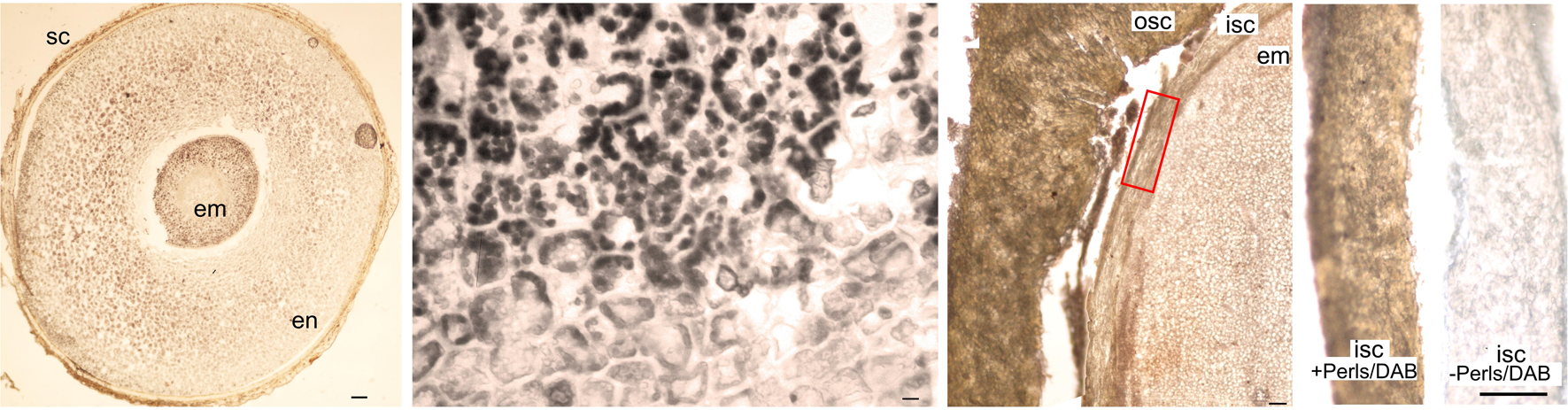
Rosid seeds that show unusual examples of storing Fe in the endosperm and in the seed coat. Fe staining patterns in *Euonymus europaeus* and *Moringa peregrina.* Seeds belonging to two different species were cut and thin cross sections were obtained. Cross sections were stained with Perls/DAB and observed under light microscope. **(A)** *Euonymus europaeus*. **(B)** a close-up to endosperm tissue of *Euonymus europaeus*. **(C)** *Moringa peregrina*, focused on a small part of the seed. Embryo, inner and outer sections of the seed coats are visible. Moringa peregrina does not possess an endosperm and has a thick two-layered seed coat. **(D)** dissected inner seed coats of *Moringa peregrina.* Left, Perls/DAB stained, right unstained control. Fe staining appeared brown in **(A, C, D)** and black in **(B)**. Seed coat in **(A)** and outer seed coat in **(C)** were already brown before the staining, thus the color does not reflect the stained Fe. In contrast, inner seed coat of *Moringa peregrina* is white, and brown color in **(C)** shows the stained Fe. sc: seed coat, en: endosperm, osc: outer seed coat, isc: inner seed coat, em: embryo. Bar is 0.1 mm for **(A)**, 0.01 mm for **(B)**, 1mm for **(C, D)**.

Finally, the distinct Fe distribution of Fe in *Euonymus europaeus* and *Moringa peregrina* seeds were further investigated by synchrotron X-ray fluorescence spectrometry. Synchrotron X-ray fluorescence confirmed the Fe accumulation in the endosperm of *Euonymus europaeus* and in the inner seed coat of *Moringa peregrina* (Fig. 7). In addition to Fe, synchrotron analysis further revealed the distribution of other metals. In *Euonymus europaeus*, Fe overlapped phosphorus (P) and Mn and was distributed through the embryo and endosperm. Zn was accumulated exclusively in the embryo.

**Figure 7:**
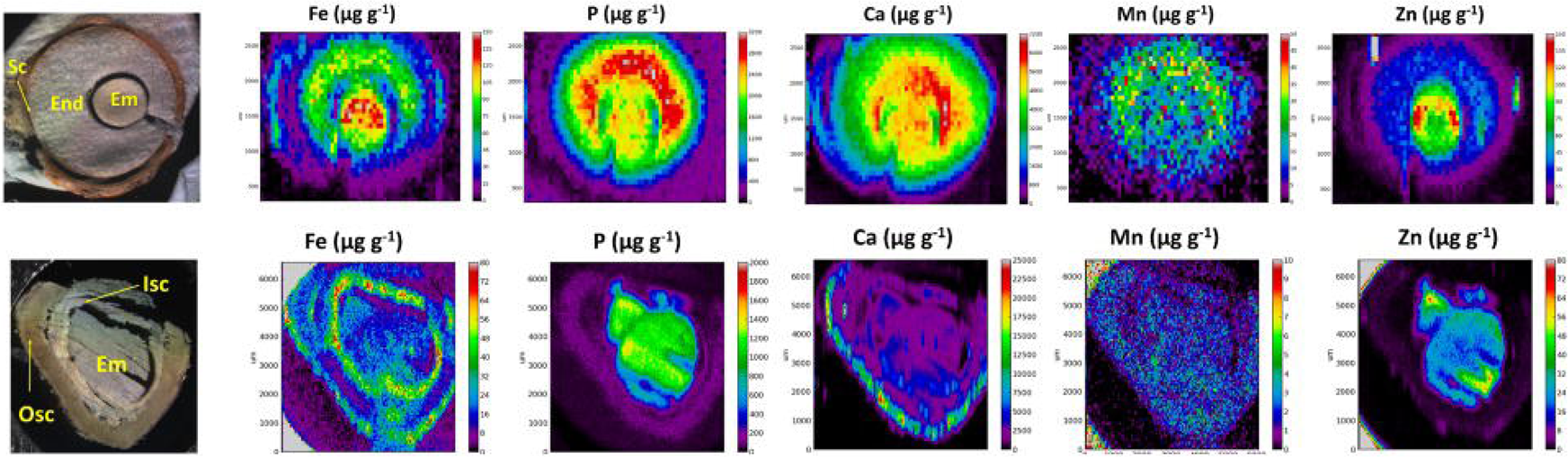
X-ray fluorescence spectroscopy confirms endosperm and seed coat of seeds can possess large Fe reserves. Spatial distribution of metals in E*uonymus europaeus (top panel)* and *Moringa peregrina* (bottom panel). The first column of images shows light microscopy images. Other columns show distribution of elements; Fe, P, Ca, Mn and Zn, respectively. Colorbar indicates concentration. sc:seed coat, en:endosperm, isc: inner seed coat, osc: outer seed coat, em: embryo.

In *Moringa peregrina*, Fe concentration was not only high in the inner seed coat as previously revealed by Perls/DAB (Fig. 6) but also in the outer seed coat (Fig. 7). Fe specifically localized to the outer side of the outer seed coat and throughout the inner seed coat. Outer seed coat of *Moringa peregrina* was also highly enriched in calcium (Ca), in a region overlapping Fe, but not other metals. In contrast to the Fe and Ca, Zn was confined to the embryo.

### Discussion

Iron deficiency is a widespread nutritional disorder worldwide. Most of the Fe in kernels of staple crops are largely unavailable to human because it is associated with antinutrients such as phytate. Directing Fe from phytate containing vacuoles to phytate-free compartments, in particular to plastids; where Fe is rather bound to ferritin, producing a more bioavailable complex, might be a beneficial strategy. This strategy requires identification of a plastid-localized strong Fe transporter. To find such transporters, using Perls/DAB staining and X-ray based methods, many seeds that belong to distinct plant lineages were screened for Fe accumulating plastids. This screening revealed that in *Carica papaya* seed, large Fe reserves exclusively localize to amyloplasts.

Among all the samples analyzed, *Carica papaya* seeds showed the most interesting clear-cut Fe hot spots. Despite the presence of a large endosperm, Fe signal appeared exclusively in the embryo (Fig. 5). Furthermore, although most of the cellular space inside the embryo was covered by large vacuoles, Fe was specifically localized in a large number of tiny vesicles like amyloplasts. Previous reports confirm that embryo axis and cotyledons of *Carica papaya* are rich for amyloplasts (dos Santos et al., 2009). To the best of our knowledge, amyloplasts have never been reported to represent the main Fe reserves in seeds of any species. For example, Cvitanich (2010) determined a large number of amyloplasts that contain ferritin in Phaseolus seeds; however, main Fe reserves localized to the cytoplasm. In papaya seed, clearcut-Fe distribution indicates a presence of a strong Fe transporter localizing to the amyloplast membrane. Since plastid Fe is generally considered as a bioavailable form of Fe, this transporter is promising in biofortification approach to increase both bioavailability and concentration of Fe at the same time. Future studies should investigate speciation of Fe in this plastid to confirm its association with ferritin protein and subsequently identify the metal transporter responsible for this plastid-localized Fe distribution.

In this study, analysis of Fe distribution in seeds also reveal the general Fe distribution patterns in species of Rosid seeds. Fe is mostly stored in the embryo part of the seeds (Fig 1-3, 5 and in all other plants that were examined in Fig. 8) and this seems to be a general trend at least in Rosids. In contrast, monocots store large concentrations of Fe in the endosperm (Lu et al., 2013; Singh et al., 2013, 2014; Vatansever et al., 2017). This difference may be explained by the decrease in endosperm size during evolution (Finch-Savage and Leubner-Metzger, 2006; Forbis et al., 2002). Early appeared species, such as monocots almost never have an embryo occupying more than half of the total seed volume. In contrast, in later appeared species (i.e., Rosids) endosperm shrinks as the seed develops and embryo occupies most of the seed. Since endosperm is a nutritive tissue, as it degrades, nutrient storage function must be taken over by the embryo itself. Therefore, we suggest metal accumulation function of the endosperm, at least for Fe, is almost completely taken over by the embryo as an evolutionary trend. In this process, metal transporters that would create sinks in the embryo might be favoured.

**Figure 8:**
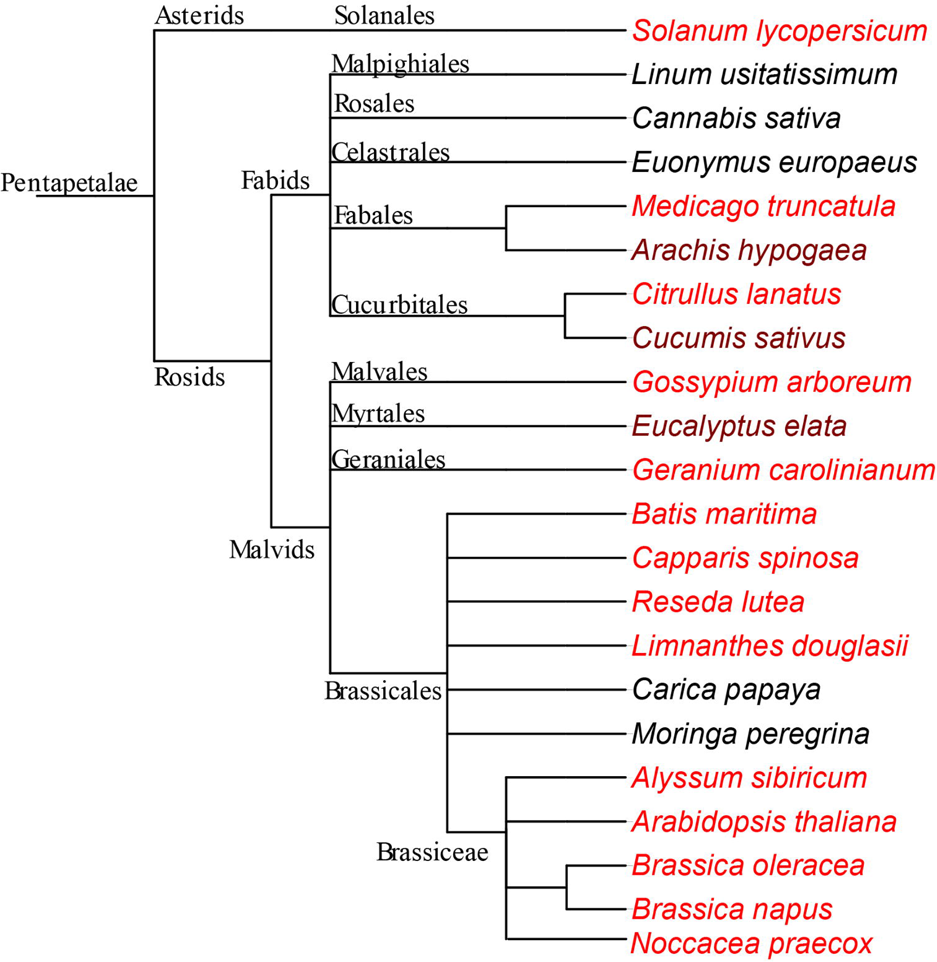
Fe enriched endodermis is a conserved feature in Rosids. Schema phylogenetically represents species that are used in this study. The phylogenetic tree was generated using NCBI taxonomy database and exported with the version 3 of the Interactive Tree of Life (iTOL) (Letunic and Bork, 2016). Both dark and light red coloured species were enriched with Fe around provascular strands. In the darker-red-coloured species, endodermis Fe does not represent the highest concentration of Fe, in contrast to the lighter-red-coloured species. The species which are not red, do not specifically accumulate large Fe reserves in their endodermis. See discussion for the interpretation of the negative data.

Besides the general trends among seeds, exceptional distribution of Fe was also detected. Embryo Fe levels could be exceeded by the endosperm or even by the seed coat (Fig 6, 7 and (Moraghan et al., 2002). In principle, the embryo is the final destination for nutrients in the seed, as the mature coat is dead while the endosperm is sterile. Although Fe that is trapped in the coat is not readily available to the germinating embryo, seed coat can contain a large amount of Fe (Fig. 7 and (Cvitanich et al., 2010; Moraghan et al., 2002)) High concentrations of trapped Fe in the seed coat can be related to the presence of chelators, such as tannins, which bind strongly to nutrients and immobilize them (Lombardi-Boccia et al., 1995). To get a better overall picture about Fe storage in the seed coats of diverse species, studies in the future should use fluorescent Fe stains as an alternative to Perls that can localize Fe despite the colored background (Park et al., 2014), where high resolution X-ray fluorescence based imaging is not a feasible option. Note that zinc is not trapped in the seed coat, which might be explained by its lower affinity to common plant chelators compared to Fe, Mn and Ca (Fig. 7). In contrast to the dead seed coat, Fe stores in the endosperm are remobilized during germination to support the germinating embryo (Becraft, 2007; Regvar et al., 2011). Monocots, including major staples such as wheat and rice, accumulate large concentrations of Fe in the endosperm, but specifically in the tiny outermost layer, aleurone. Aleurone remains alive in the mature seed, in contrast to starchy endosperm cells which are almost devoid of Fe (Lu et al., 2013; Singh et al., 2013, 2014; Vatansever et al., 2017). In Rosids, endosperm rarely stores a significant amount of Fe (only in a few species, see Fig. 6A, B, Fig. 7 and Supp. Fig.4). Interestingly, X-ray fluorescence showed Fe in the endosperm colocalizes with phosphorus (P) (Fig. 7A). Localization of P generally mirrors phytate distribution in seeds, indicating Fe was trapped by phytate in the endodermis before it could reach to the embryo (Iwai et al., 2012). In summary, the principal storage organ for Fe is the embryo in Rosid seeds; however, Fe reserves can be restrained in the way to the embryo, probably due to the immobilization by chelation in the endodermis or in the seed coat.

We showed that Fe accumulation exhibits hot spots in embryos, which correspond to the endodermis (Fig. 8). Among other Fe enriched regions, endodermis was detected as the only conserved one among distinct plant lineages (Fig. 8). In *Arabidopsis thaliana*, Fe enriched endodermis is strictly dependent on vacuolar Fe transporter VIT1 (Kim et al., 2006). Therefore, we assumed Fe-enriched endodermis in species other than *Arabidopsis thaliana* was also due to the presence of VIT1 homologs in those species. In few species, Fe enriched cells go beyond the single endodermal cell layer to nearby cortex cells (compare Fig. 2B with 2D, also see Ibeas et al. (2017)). Therefore, the question arises whether VIT1 can localize to cells other than endodermis or is it strictly cell-type specific. As elegantly shown by Roschzttardtz et al. (2009) using *shr* mutant lacking the endodermis, VIT1 does localize to cortex cells if endodermis does not exist. This may indicate regardless of whether it is restricted to a single cell layer or not, the typical ring-shaped Fe localization around provascular strands is always due to the expression of *VIT1*.

Although VIT1-mediated Fe enrichment was conserved even in distinct orders, few species did not exhibit this phenotype (Fig. 5-8). Among those seeds, endodermis in papaya embryo was devoid of any staining, while, in all the rest, staining in endodermis were not noticeably higher. This may pose the question whether VIT1 has been lost in these species during the course of evolution. However, conservation of a VIT1 domain (PF01988) (Fig. 4A and Supp. Fig. 2, 3) and distribution of these species with the ones that showed Fe enriched endodermis in the phylogenetic tree (Fig. 8) do not support this possibility. Failure to detect a VIT1-mediated Fe-enriched endodermis was not attributed to the absence of a functional VIT1 but to the presence of other transporters which take over VIT1’s function in storing Fe. Studies with loss of function mutants (Eroglu et al., 2017; Kim et al., 2006) indicates that Fe accumulation patterns are eventually determined by the single dominant transporter. For example, VIT1’s presence prevents MTP8’s impact on Fe storage in *Arabidopsis thaliana*. Likewise, when a more preferential Fe transporter is present, VIT1’s impact on Fe distribution (i.e., Fe-enriched endodermis) might become insignificant. In addition to this model, the imaging techniques used in the study are technically limited in revealing a less pronounced Fe store in the presence of a highly pronounced one. For instance, in *Arabidopsis thaliana*, endodermal cells produce the only significant Fe signals by X-ray analysis (Kim et al., 2006) or Perls/DAB staining (Fig. 1A) and the signal from the rest of the cells is unnoticeable. Nevertheless, the rest of the cells in the embryo account for half of the total Fe in the seed, as shown by precise quantitative techniques (Ramos et al., 2013). Taken together, VIT1 gene and its associated phenotype is well conserved despite some seeds show alternative Fe hotspots.

Rosids is a huge lineage including 80000 species belonging to 147 families, making more than a third of all angiosperms (Hedges and Kumar, 2009; Soltis, 2005). The current study shows seeds store Fe mainly in embryo in Rosid species. This Fe is not equally distributed but most often concentrated in the innermost cell layers, endodermis and sometimes cortex, in a VIT1-dependent manner. Future studies should carefully examine beyond Rosids to pinpoint in which stage of plant evolution Fe enriched endodermis appeared as a new feature. Furthermore, in contrast to most of the other Rosid species, *Carica papaya* embryos diverted most of its Fe reserves into the plastids. This indicates the presence of a strong Fe transporter on the plastid membrane. Once identified, such a transporter can become a promising tool for future biofortification attempts, to divert seed Fe into compartments that are free from antinutrients.

## Acknowledgments

Thanks to Asci Murat Mihladiz (Capparis Research&Development Center, Burdur, Turkey), Dr. Emre Cilden (Hacettepe University, Ankara, Turkey) for sharing seed stocks with us. We are grateful to Dr. Ali Veral and his team, especially Ebru Çanlı (Ege University Hospital, Izmir, Turkey) for excellent technical help in preparation of cuttings. Thanks to Recep Vatansever and Ferhat Celep for fruitful discussions. Authors also acknowledge Primoz Pelicon and Primoz Vavpetic (JSI) for the help with PIXE analysis and Iva Bozicevic Mihalic (Elettra Sincrotrone Trieste) for the help with the synchrotron measurements. Elettra is acknowledged for the provision of the beamtime (project 20175078).

## Author contributions

SE and NK collected the seeds and performed histochemistry; KM and AK, X-ray analyses; EF, bioinformatics. SE and BT conceived and planned the project. SE wrote the article with contributions of all the authors.

