## supplementary file for "VIT1-dependent Fe distribution in seeds is conserved in dicots"

**Supp. Fig. 1.**
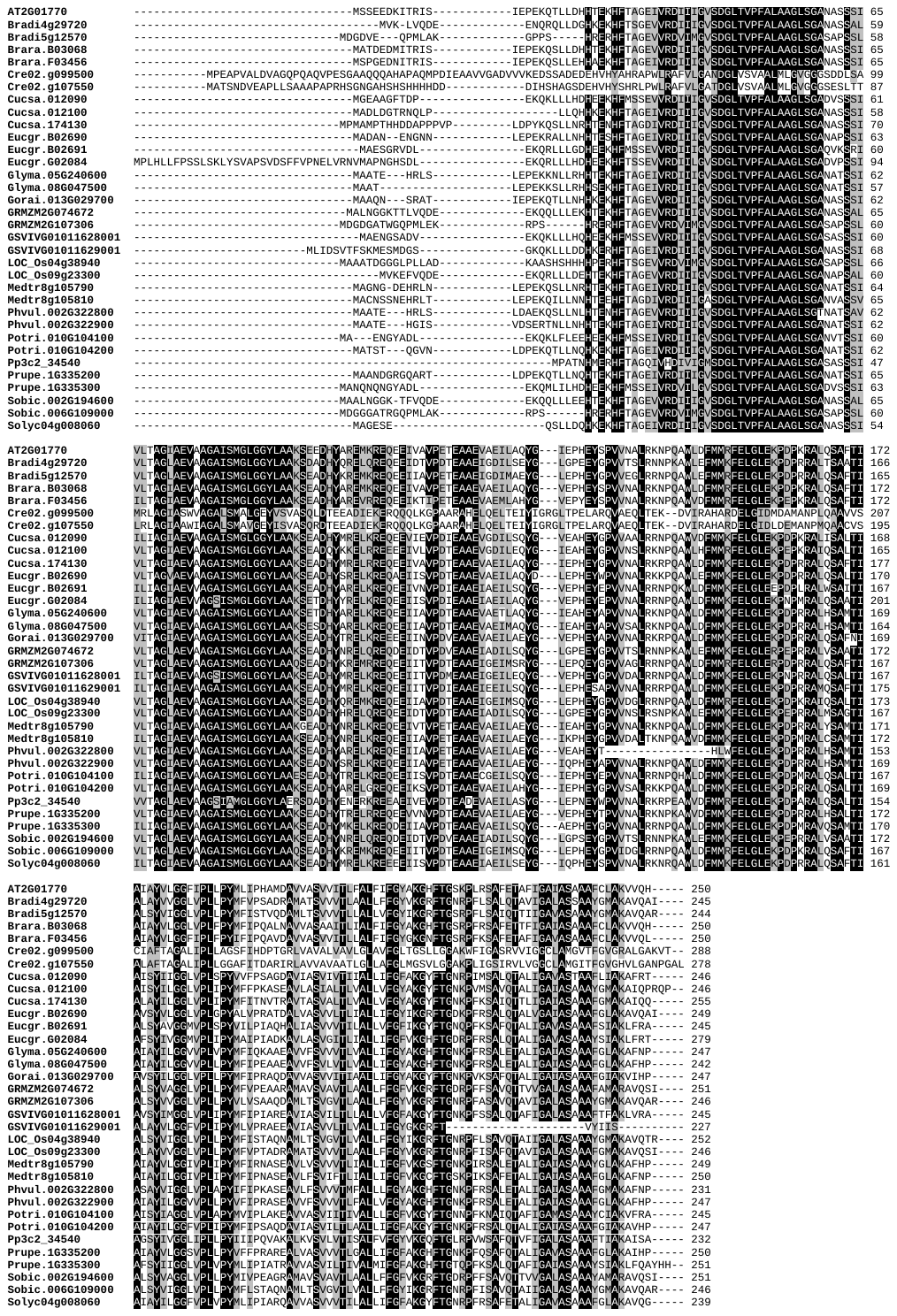
 Alignments of 34 VIT1 sequences from 18 different plant species.

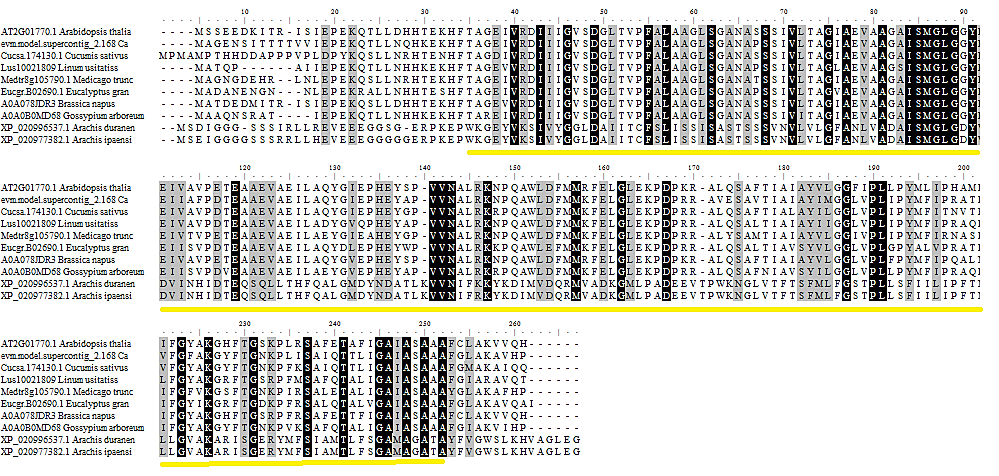

**Supp. Fig. 2.** Alignment of VIT1 from species which were chosen for seed Fe staining. The yellow line shows the conserved VIT1 (PF01988) domain structure

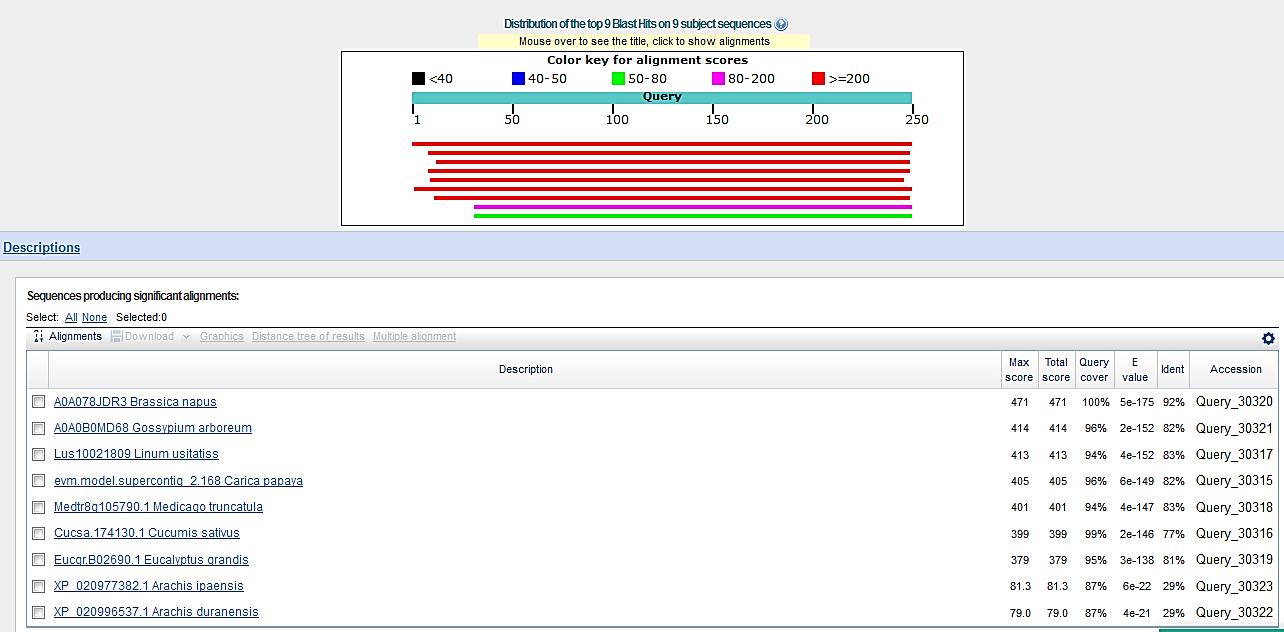

**Supp. Fig. 3.** Comparison of *A. thaliana* with the other nine plant VIT1 sequences using protein blast.

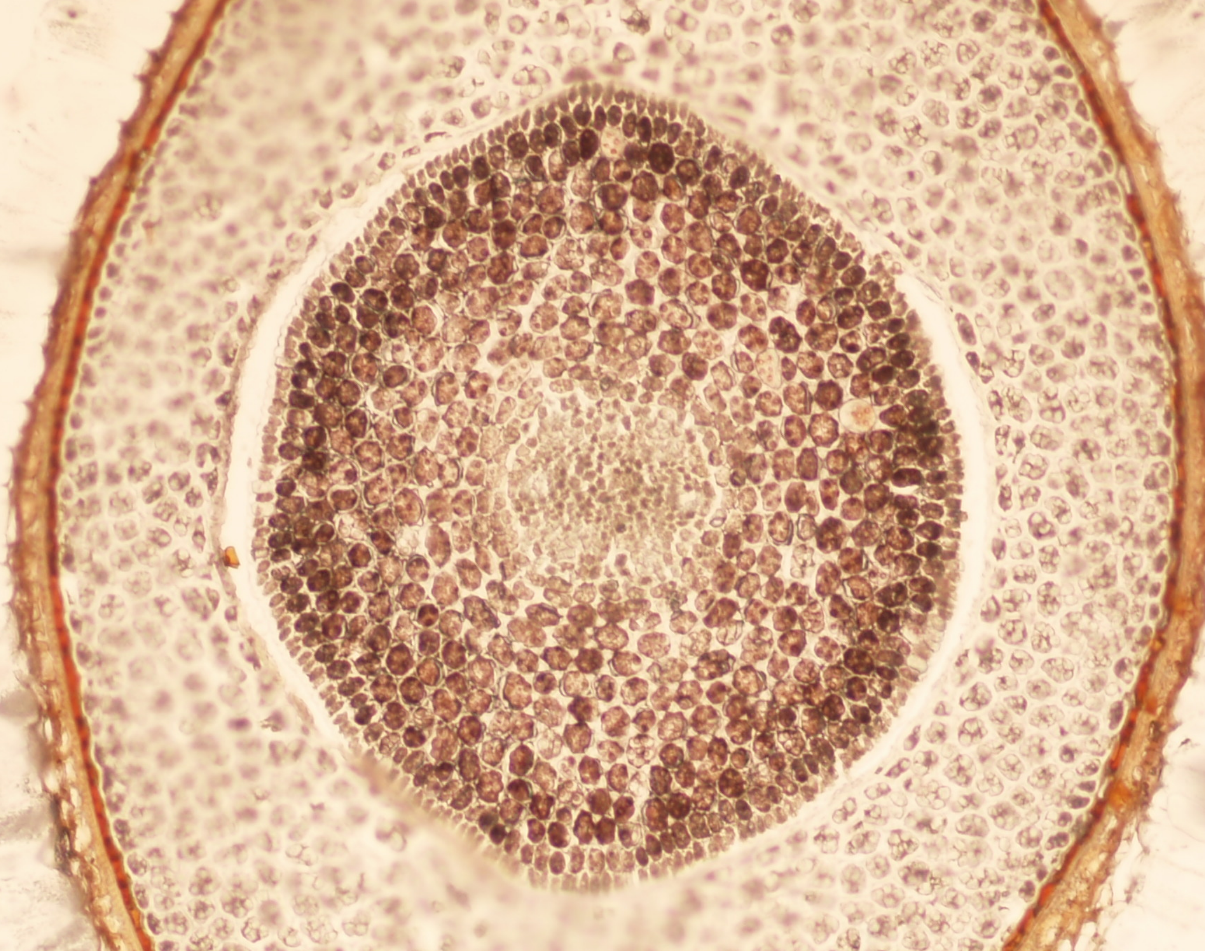

**Supp. Fig. 4.** Perls/DAB staining of common flax (*Linum usiatiss*).

**Supp. Table 1.** Biochemical properties of VIT1s according to their protein sequence analyses.

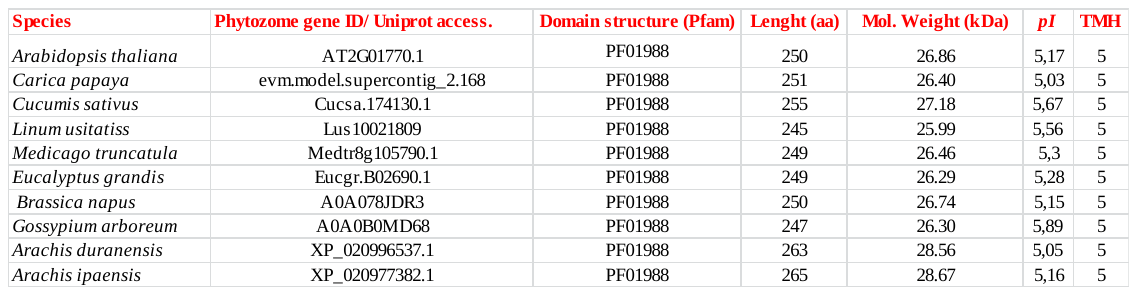
